# On the reliability of wearable technology: A tutorial on measuring heart rate and heart rate variability in the wild

**DOI:** 10.1101/2022.10.02.510535

**Authors:** Veronica Dudarev, Oswald Barral, Chuxuan Zhang, Guy Davis, James T. Enns

**Author notes:** **Author Note** Data and R code generated for this tutorial are available at https://zenodo.org/badge/latestdoi/520639317. VD’s involvement in this research was funded by Mitacs Accelerate Postdoctoral fellowship (IT23473). The research costs associated with this study were funded by a Discovery Grant to JTE from the Natural Sciences and Engineering Council of Canada and by HealthQB Technologies, Inc. GD is the founder and CEO of HealthQb Technologies, Inc., which supported the study by providing salary to CZ and OB, measurement equipment, and covering participant reimbursement. HealthQb Technologies was not involved in either designing the study, or data analyses and interpretation. Correspondence should be addressed to Veronica Dudarev, 2136 West Mall, Vancouver, B.C. Canada, V6T 1Z4.

## Abstract

Wearable sensors are quickly making their way into psychophysiological research, as they allow collecting longitudinal and ecologically valid data. The present tutorial considers fidelity of physiological measurement with wearable sensors, focusing on reliability. We elaborate why ensuring reliability for wearables is important and offer statistical tools for assessing wearable reliability for between participants and within-participant designs. The framework offered here is illustrated using several brands of commercially available heart rate sensors. Our hope is that by systematically quantifying measurement reliability, researchers will be able to make informed choices about specific wearable devices and measurement procedures that meet their research goals.

## Introduction

A bathroom scale is a reliable measure of one’s weight, provided one stands still on the scale for several moments. Yet one is likely to discard the measurement shown by the scale if one is startled by a spider during these moments. Trying to measure cardiac signals with a wearable sensor is a little like trying to measure one’s weight while dancing on the scale. The fidelity of the measurement will depend not only on sensor’s accuracy but also on the environmental conditions under which the measurement was taken.

The recent rapid proliferation of wearable sensing technology has been accompanied by many tests of their validity (Barrios et al., 2019; Bent et al., 2020; Dur et al., 2018; Hernando et al., 2018; Kinnunen et al., 2020; Koskimäki et al., 2018; Menghini et al., 2019; Steinberg, Yuceege, Mutlu, Korkmaz, Van Mourik, et al., 2017), usually by examining the correlation of the wearable’s signal in a quiet environment with that of trusted laboratory equipment (van Lier et al., 2020). What is usually overlooked in these tests is that most laboratory measurement procedures severely limit participants’ bodily movements and cognitive activities (Berntson et al., 2007). In sharp contrast, wearable devices promise to measure the same physiological signals across a wide variety of environments and bodily states. Yet without further testing, there is no guarantee that wearables will yield accurate measurements in all contexts.

For example, Empatica’s electrodermal activity (EDA) measurement showed high agreement with the EDA measurement taken under laboratory conditions. Yet in a study that measured EDA with Empatica for 20 hrs per participant in their daily lives, 78% of the measurements were artifacts and no meaningful analysis could be performed with the remaining data (Zheng & Poon, 2016). In another study, which aimed to establish the validity of Empatica’s HRV measurement against a Holter ECG monitor in 24-hour ambulatory monitoring, the reported reliability of Empatica’s measurement of heart rate variability (HRV) was lower than that of Holter device, and the proportion of missing data was higher (Van Voorhees et al., 2022).

The present paper focuses on the question: *when* can measurement from a wearable device be trusted? Keeping in mind that the appeal of wearable sensors is to measure physiology under conditions where a benchmark device (e.g., ECG) is not viable, we offer several statistical tools that allow assessing measurement fidelity *without referencing a second device*. The tools we offer are based on the concept of reliability (Revelle & Condon, 2019).

It is textbook knowledge that measurement fidelity can be decomposed into two components: validity and reliability. Validity denotes measurement *accuracy* – usually determined as correspondence of measurement to another gold-standard measurement of the same variable. Reliability refers to measurement *precision* – that is, consistency of several measurements taken in the same conditions and/or with the same equipment. It is useful to distinguish between validity and reliability, as can be demonstrated by considering an example of drunk dart thrower. What does being drunk do: limit accuracy, limit precision, or does it limit both? As shown in Figure 1 (https://conjointly.com/kb/reliability-and-validity/), the two are orthogonal. If a drunk lacks accuracy, they behave as in A, being consistently off the mark but with reasonable precision. If they lack precision, they do most poorly in B, though their average accuracy is still good. If they lack both, it is box C. This example helps understand why low reliability makes accuracy hard to determine. There is too much scatter in the data to estimate the mean with confidence. It also helps understand how high reliability does not guarantee accuracy: the dart thrower may be very precise (reliable) but consistently off the mark.

**Figure 1.**
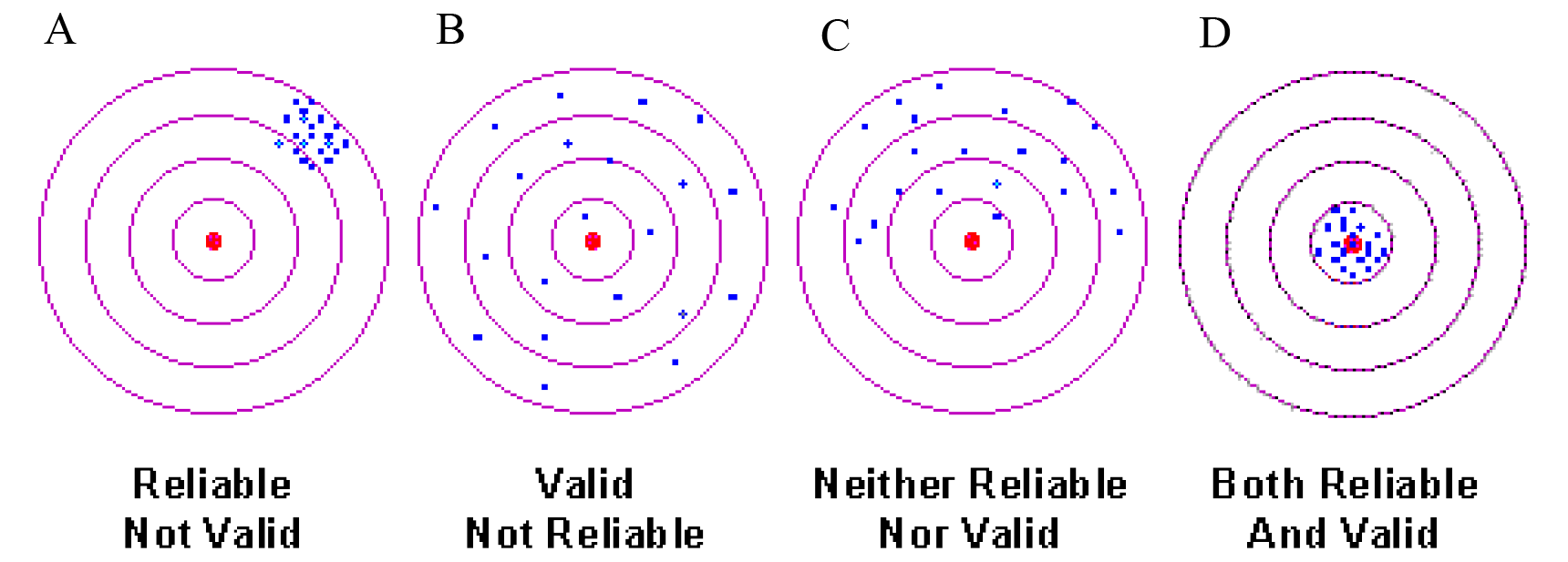
Illustration of the relationships between validity and reliability using the example of drunk dart thrower (image taken from https://conjointly.com/kb/reliability-and-validity/).

We begin by noting that the theory of measurement reliability was originally developed for assessing fidelity of subjective reports (questionnaires, ratings); not for physiological measures. When participants are responding to questionnaires, one cannot fully control the conditions under which they are being filled out (noise, distractions, time pressure) and what factors other than the question may be influencing their responses. Tools to assess response reliability were designed to measure this uncertainty. Psychophysiology borrowed these insights from the theory of reliability to ensure that psychophysiological measurements and associated laboratory protocols resulted in consistent and reliable estimates (Berntson et al., 2007; Di Nocera, Ferlazzo, Borghi, 2001; Tomarken, 1995). Here we do the same for signals from wearable sensors, while taking into account recent developments in reliability theory itself (Revelle & Condon, 2019). Notice that the reliability analyses we propose are performed on data readily available from any wearable device, without additional devices and measurements.

To the best of our knowledge, wearable device reliability has not been considered in detail before. Kleckner et al. (Kleckner et al., 2021) mentioned that for a wearable sensor to be accurate its measurement has to be reliable, but their proposed framework for choosing a wearable device for research offers no guidelines on assessing its reliability. Here we fill this gap, by offering several readily accessible tools to estimate reliability of measurement with respect to a particular goal.

### Definitions

Measurement reliability refers to the consistency of the estimates obtained under hypothetically equivalent conditions; the complement of reliability is measurement uncertainty. Not only does this uncertainty undermine the accurate measurement of a physiological state, but it seriously weakens the ability to use the measured physiological state as a predictor of other outcomes. Spearman (Spearman, 1904) noted this issue long ago, showing that low reliability in a predictor variable directly decreases the measured agreement between predictor and outcome variables (Johnson, 1944; Nimon et al., 2012; Revelle & Condon, 2019). Despite this well-known relationship between measurement error and measures of association, many statistical treatments assume that predictor variables (e.g., predictors in a regression) are measured without error, simply because accurate predictor data is the standard assumption in the application of linear regression (Sklar et al., 2021). It is also the case that low predictor reliability weakens the results of many statistical tests other than correlations (Nimon et al., 2012). Although there are ways to assess reliability (Revelle & Condon, 2019) and to correct some statistics for low reliability (Spearman, 1904), many researchers do not use them (Nimon et al., 2012; Revelle & Condon, 2019). To summarize, the low reliability of a physiological measurement can prevent one from discovering its true association with other variables.

It is also important to note that not all measurement variability is measurement error. Rather, researchers try to distinguish among sources of potential variance in a measurement, and accordingly, the consequences of these various types of variability on measures of reliability (Nimon et al., 2012; Revelle & Condon, 2019). Human physiology is affected by at least two broad groups of factors: constitutional factors and situational variables (Fatisson et al., 2016; Shaffer & Ginsberg, 2017). Depending on the aims of a study, either of these can be considered as noise. For instance, in a study investigating stable differences between individuals, such as differences in personality or physiological traits, situational and state differences between people are a source of noise. Conversely, in studies comparing situational differences, individual differences in personality or physiological traits are a source of noise (Hedge, Powell & Sumner, 2018). Therefore, the type of reliability to be considered depends on the goals of the study.

Constitutional factors refer to enduring or trait-like states of the body. In the case of cardiac measurements, these factors can be intuitively linked to gender, age, body mass index, physical fitness, and chronic medical conditions. This assembly of stable personal traits has predictable influences on blood pressure (Printz & Jaworski, 2004), heart rate (Sommerfeldt et al., 2019; Zhang et al., 2016), and heart rate variability (Laborde et al., 2017; Natarajan et al., 2020; Shaffer & Ginsberg, 2017). The stability of a person’s physiological parameter measured in different situations is thus referred to as **between-person reliability** (sometimes also as “relative reliability”, or parameter level measurement precision– see van Lier et al., 2020), because it indexes the extent to which a person’s parameter is stable relative to other people in different situations. For example, if one’s heart rate (HR) or heart rate variability (HRV) is generally high when compared to other people in a laboratory testing, then we would expect their HRV assessed by a wearable device to also show that it was generally higher than other people’s data measured by the same device.

The second source of cardiac variability – situational factors – refer to physical activity, stress level, and other more transient physiological states. For example elevated heart rate along with reduced HRV is associated with fever (Karjalainen & Viitasalo, 1986), inflammation (Williams et al., 2019), and acute pain (Chowdhury et al., 2021; Kasaeyan Naeini et al., 2021; Koenig et al., 2014; Lim et al., 2019), as well as mental stress (Brosschot et al., 2007; Hernando et al., 2018; Hovsepian et al., 2015) and physical effort (Perini & Veicsteinas, 2003; Tulppo et al., 1998). It is these situational factors that are spurring much of the current interest in wearable devices. The hope is that tracking users’ heart-rate biometrics will provide a useful clue for ensuring their health and wellbeing. For instance, studies that compared physically fit people to those who do not exercise as much (between-participant design) tend to show that physical fitness is associated with higher HRV (Buchheit et al., 2005; Rennie et al., 2003; Tulppo et al., 1998). It is plausible to assume then that increasing one’s fitness will result in an increase in HRV, when compared to the same person’s HRV before training. But confirming this conclusion really calls for a within-person study design – i.e., measuring HRV in the same person before and after a change in fitness (Janse van Rensburg et al., 2012; Routledge et al., 2010). A within-person comparison of biomarkers therefore calls for the assessment of **within-person reliability.** This is also sometimes referred to as “absolute reliability,” and it quantifies the stability of a sensor’s readings from the same person in a given situation/state, compared to other states of the same person (for a greater in-depth discussion of between and within-person reliability see Revelle & Condon, 2019).

### Measuring between-person reliability

Between-person reliability refers to the stability of measurement for the same person relative to other people, across time and contexts. In simple words, it is the agreement between measurements taken at different times for the same person (see Figure 2 for illustration). Let us consider a single measurement as

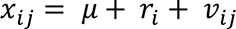

where

*x_ij_* is a measure taken from individual *i* in situation *j*.
*μ* is the population average,
*μ* + *r_i_* is the average for each participant *i*,

*v_ij_* is measurement error

Using these terms, between participant reliability can be expressed as

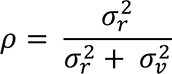

This is the traditional formulation of population Intra-class correlation (ICC - (Bartko, 1966)). For a specific sample, in its most general form, it can be computed as:

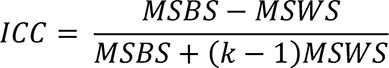

where

*MSBS* is the mean sum of squared deviations between the participants
*MSWS* is the mean sum of squared deviations within the participants
*k* is the number of measurements (which is required to be equal across participants).

**Figure 2.**
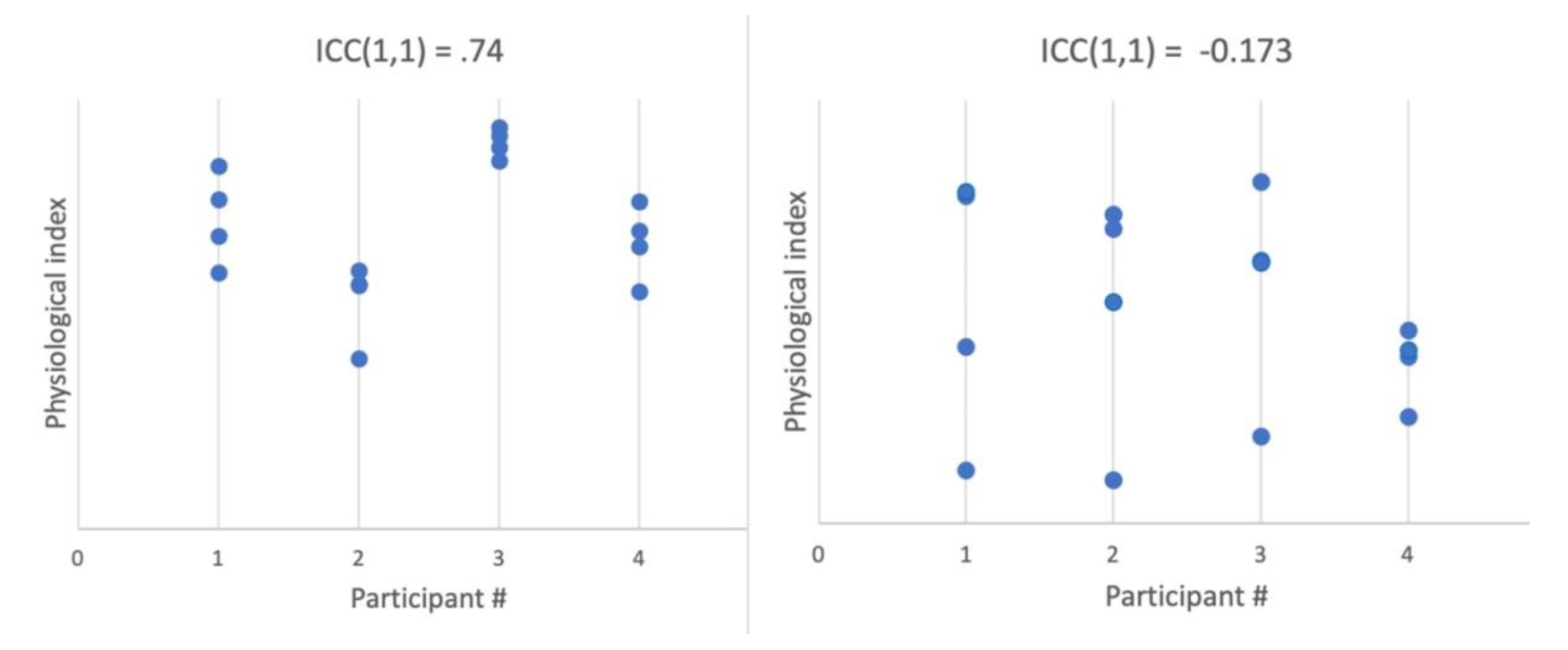
Simulated data of 4 measurements per each of 4 participants. Right panel shows an example of low discriminability between participants, i.e. low one-way single-measure ICC. Left panel shows a case in which each participant’s physiological index is highly individual, resulting in high ICC and high between-participant reliability.

Theoretically, ICC varies from −1 to 1^1^, and should be interpreted in the same way as correlation coefficient: the closer to 1 the higher the agreement, with ICC > .75 representing excellent reliability (Cicchetti, 1994).

Several different forms of ICC are available when modeling additional sources of variance. For example, consider a number of patients being examined by several doctors (raters), which is a classic case for ICC application. In this case, it is standard to model the variation among the raters using two-way ICC, based on the assumption that different people may provide ratings that are systematically different from one another (specific to each rater). For a physiological measure, when there is no a-priori reason to assume that individual sensors would be systematically different in their measurement, the most general one-way ICC would be sufficient. The formulas above represent one-way ICC, capturing between-participant variance (MSBS) against noise variance (MSWS) only. If one wishes to model individual variation along with systematic difference between measurement devices (e.g., different models) or circumstances (e.g., exercise, stress, rest), then a two-way ICC may be applicable.

In addition to deciding whether one-way or two-way ICC is most appropriate, a decision also needs to be made whether to use single measurement or multiple measurements options. Single measurement ICC estimates representativeness of a single measurement for the person’s parameter value. Multiple measurements ICC estimates representativeness of the average across all measurements. For additional details and guidelines see (Liljequist et al., 2019; Revelle & Condon, 2019).

### Measuring within-person reliability

Within-person reliability refers to the stability of a measurement taken in the same situation, for a given person. In the extreme, it would refer to two identical devices producing equal measurements when used simultaneously on the same person. For a single device, considering measurement as

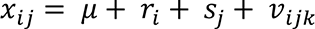

where

*x_ij_* is a measure taken from individual *i* in situation *j*.
*μ* is the population average,
*μ* + *r_i_* is the average for individual *i*,
*μ* + *s_j_* is the average for a situation *j*, and
*v_ijk_* represents the difference between multiple measurements taken for individual *i* during situation *j*.

Here, within-participant reliability would be the opposite of the magnitude of *v_jk_*. The conceptual distinction between signal and noise is crucial when considering within-person reliability and will depend on a particular design and aims of the measurement situation (for further guidance see (Revelle & Condon, 2019)).

Let us consider a simple example. Imagine we are measuring the heart rate of a particular person, taking several measurements during rest, a few more measurements during mentally stressful activity, and a few more during physical exercise. We would expect high agreement within rest measurements, and distinct but clustered measurements during each type of activity. Mixed model regression predicting heart rate measured at time 1 from heart rate measured at time 2, with situation (rest, mental activity, physical activity, recovery) as the random factor takes the following form:

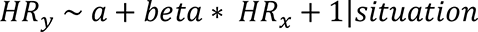

As in general linear regression approach, *beta* is the estimate of the association between the predictor and the predicted variable. In this case, *beta* quantifies the amount of agreement between measurements taken during each situation. If we test more than one person, we should add a random factor of participant, to remove variance associated with individual differences and focus on within-participant consistency:

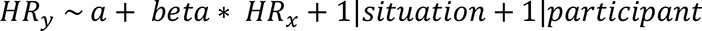

Figure 3A shows an example data simulating 4 participants in 4 types of situations where both between-and within-participant reliability (consistency) are high.

**Figure 3.**
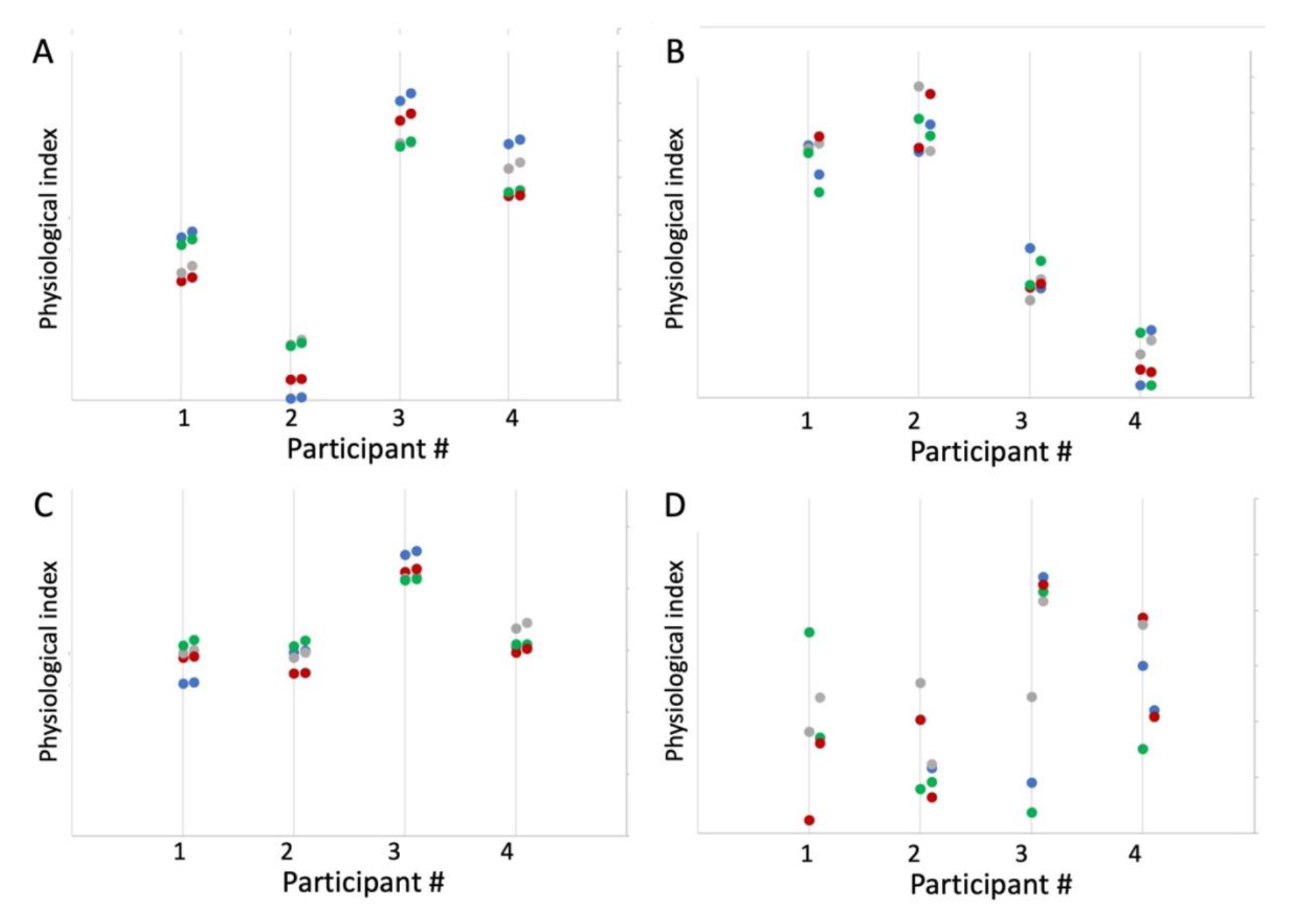
Simulated data of 8 measurements for each of 4 participants, in 4 situations (colour coded). **Panel A** shows an example of high between-and within-participant reliability, where the observations are consistent per participant and per situation. **Panel B** shows example of high between-participant reliability yet low within-participant reliability. **Panel C** shows high within-and low between-participant reliability, where datapoints are consistent within a situation yet do not reliably distinguish between different individuals. **Panel D** shows low within-and between-participant reliability.

As can be seen, this type of analysis requires two measurements for the physiological index of interest in each situation. Traditionally, with subjective responses to questionnaires, the questions were divided into two subsamples by the order of their appearance, taking either first and second half of the questionnaire as the two subsamples, or odd vs. even questions (so called split-half approach, (Spearman, 1904; Van Norman & Parker, 2018). Unlike questionnaires, physiological measurement, especially those obtained with a wearable device in ecological settings, is performed over a longer time span than is required to fill out a questionnaire, often with the aim of quantifying a change in the person’s state from one measurement to the next. Given the volatility of physiological measurement in time, the closer two samples occur in time, the more similar we would expect them to be. In other words, it is reasonable to expect that time is a systematic factor that must be taken into account. Assigning each datapoint a number by order of its acquisition, then aggregating (averaging) all odd and even datapoints allows one to measure consistency between instances that are taken **as close in time as possible**. We will refer to this as time-sensitive sampling.

An alternative approach that is not time-sensitive might involve dividing all measurements into two subsamples randomly. With just one instance of such division, there is a non-zero chance that by coincidence the two subsamples will be uncharacteristically similar or dissimilar. However, if a random split is performed multiple times, we can estimate within-participant reliability from the resulting distribution of *betas*. In the next section, we will compare these two methods of dividing the datapoints to provide an assessment of within-person reliability.

### Empirical examples

We first apply the approach proposed here to a case of a commercially available PPG-based sensors of cardiac biometrics (heart rate and heart rate variability). The main aim is to demonstrate the degree to which the measurement reliability of a wearable sensor varies with the conditions of everyday life. We focus on the most discussed factors that affect measurement fidelity of wearable sensors: (1) the make of the sensor (hardware + software) and (2) physical activity of the user (Barrios et al., 2019; Thomson et al., 2019). We do this by analyzing a publicly available dataset that contains heart rate recorded by 6 commercially available PPG-based sensors (Bent & Dunn, 2021). We then move to apply the framework proposed here to data collected outside of the laboratory, where comparison to benchmark ECG is not viable. The naturalistic data were collected from 10 healthy participants who were wearing another commercially available PPG sensor, Biostrap, for a week. We compare the reliability of data acquired during sleep and during active wakefulness. In addition, to demonstrate the relationship between the reliability of a measurement and the magnitude of its correlation with another variable (Revelle & Condon, 2019; Spearman, 1904) we test the correlation between the two biometrics explored here with a measure of participants’ mood.

## Study 1: Reliability of 6 wearable sensors of cardiac biometrics

Bent et al (Bent et al., 2020) conducted a study comparing 6 different wearable devices against ECG to determine measurement fidelity of these devices under different conditions. The data (beats per minute from each device) are publicly available (Bent & Dunn, 2021). Here we analyze this dataset by applying the estimation procedures already described to assess the measurement reliability of the devices *without referencing ECG*. We compute between-and within-participant reliability for (a) the 6 wearable devices, and (b) different activities.

### Method

A total of 53 participants were tested with 6 wearable sensors while engaging in 4 types of activities: rest, paced breathing, physical activity (walking), and typing. In between these activities, the sensor was on the participant’s wrist and still recording, and so we include the rest period as the 5^h^ type of activity: task transition. Participants were wearing one or two devices at a time, repeating the activities several times.

### Results

#### Data processing

Heart rate was measured as beats per minute (BPM). Of these, we removed all the 0 values, and then values more than 2 standard deviations above or below each participant’s average. For between-participant reliability, BPM datapoints were averaged per device and activity type for each participant.

#### Between participant reliability

A one-way random single-measure ICC(1,1) was computed for the 44-53 participants’ mean BPM, separately for each device, with the 5 activity types as the different measurement instances. Table 1 and Figure 4 show the results. The 6 devices appear to have unequal between-participant reliability, Biovotion showing the highest, and generally good reliability of .65 (but also the largest number of participants without data), Empatica and Miband showing only fair reliability of .38 and .37 respectively, and the other three devices showing good reliability (Cicchetti, 1994).

**Figure 4.**
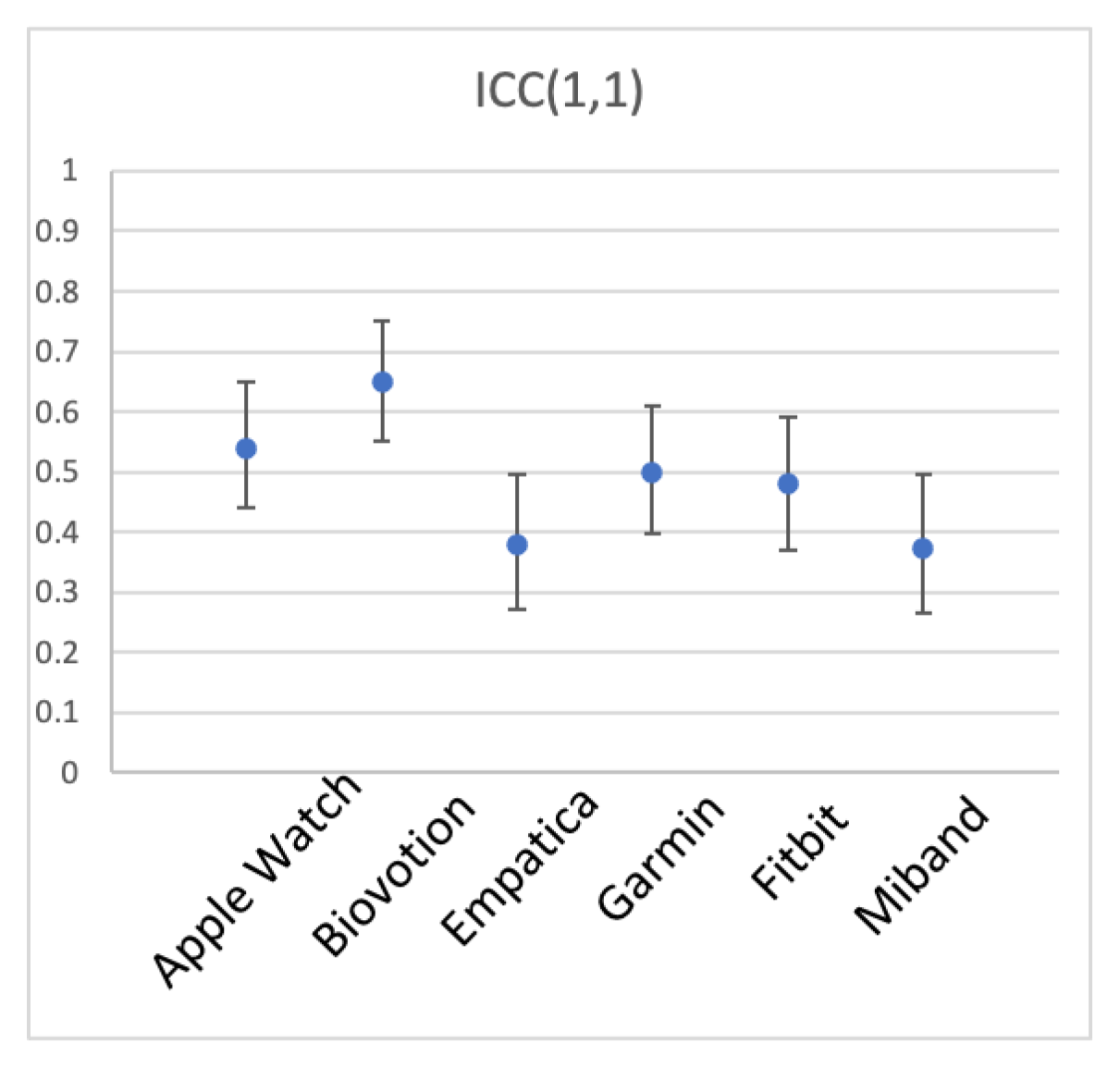
Between-participant reliability of HR for the 6 brands of wearables devices. ICC(1,1) is shown, error bars represent 95% CI.

**Table 1.** *Between participant reliability of HR for the 6 wearable sensors, listed in alphabetic order. Number of participants (N) varies per sensor because of missing data*.

| Device | ICC(1,1) | 95% CI | N |
| --- | --- | --- | --- |
| Apple Watch | .54 | [.44 .65] | 53 |
| Biovotion | .65 | [.55 .75] | 45 |
| Empatica | .38 | [.27 .496] | 53 |
| Garmin | .50 | [.397 .61] | 52 |
| Fitbit | .48 | [.37 .59] | 53 |
| Miband | .37 | [.27 .496] | 50 |

We then explored whether different conditions of measurement – in this case, different activities – produce data that is more or less representative of individual participant’s heart rate (between participant reliability). To this end, we computed ICC(1,1) for the 5 types of activity, with devices serving as measurement instances. Table 2 shows the results. Breathing, transitioning between activities, and rest elicited the most reliable measurements across devices (ICC(1,1) of .66, .597, and .54, respectively). Physical activity (walking) and typing produced ICC(1,1) of just .37, suggesting that measurement was quite noisy during these activities.

**Figure 5.**
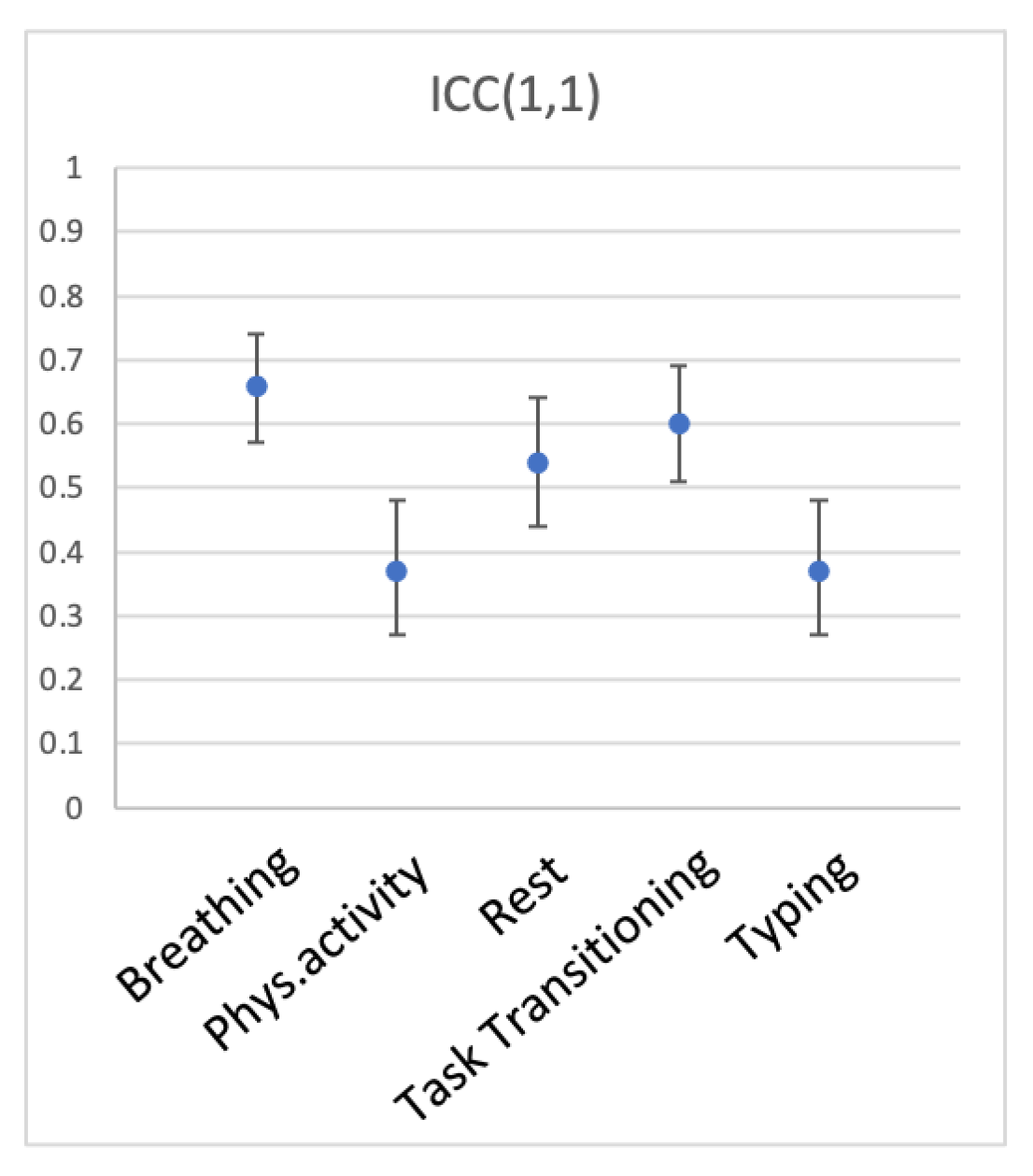
Between-participant reliability of HR across wearable devices for the 5 types of activity. ICC(1,1) is shown, error bars represent 95% CI.

**Table 2.** *Between participant reliability of HR for the 5 activities, listed in alphabetic order*.

| **Activity** | **ICC(1,1)** | **95% CI** | **N** |
| --- | --- | --- | --- |
| Breathing | .66 | [.57 .74] | 53 |
| Physical activity | .37 | [.27 .48] | 53 |
| Rest | .54 | [.44 .64] | 53 |
| Task Transition | .60 | [.51 .69] | 53 |
| Typing | .37 | [.27 .48] | 53 |

### Interim Discussion

Between participant reliability was examined in the dataset containing heart rate of 53 participants measured with 6 devices during 5 types of activities. Reliability varied between the 6 devices, with Biovotion and Apple Watch showing highest reliability, closely followed by Garmin and Fitbit, with Empatica and Miband showing lower reliability. Interestingly, comparison to ECG measurement reported in Bent et al. (Bent et al., 2020) revealed that the deviation from ECG was lowest for Apple Watch, followed by Garmin and Fitbit, followed by Empatica and Miband, followed by Biovotion. That is, measurement fidelity as assessed by comparison to ECG (validity) and as assessed by between participant reliability of the measurement itself (reliability) match closely, with Biovotion being the only exception. It is textbook knowledge that reliability is a necessary, but not sufficient condition for validity of measurement. And this is exactly what the data shows for Apple Watch and Miband. As the reliability of wearable devices decreases across brands, their validity decreases as well. Note too that Biovotion is a clear example of a device that is reliable (not much internal noise), yet not valid (does not correspond to a benchmark device). Thus, high reliability does not guarantee high validity (see Figure 1). But it is also true that low reliability makes it difficult to determine validity at all. However, once validity is established, is it possible that measurements are unreliable? We addressed this question by investigating different participant activities.

Participants’ activity generally affected measurement reliability as expected, with calmer states (breathing, transitioning, rest) producing higher reliability than more intense activities (walking, typing). This is consistent with multiple previous studies, including Bent et al. (2020), who reported higher reliability for measurements taken during rest, and reduced reliability with increased levels of activity.

#### Within-participant reliability

Within-participant reliability was assessed using a split-half approach and mixed model regression. We could not use a time-sensitive approach because time stamps were not available in the dataset; we therefore used a random split-half approach. To explore within-participant reliability of the 6 devices, we split each participant’s heart rate datapoints during each activity into random halves 1000 times, computing mixed-model regression with participant as the random factor each time. Reliability was estimated as the average beta across the 1000 iterations. To explore the reliability of measurement during different activities, we split each participant’s heart rate as measured with each device into random halves 1000 times, submitting that to mixed-model regression every time.

Figure 6 and Table 3 show within-participant reliability for the 6 devices. As can be seen from the table, all devices had excellent reliability in measuring heart rate across different situations within participant. It can also be seen that reliability of Fitbit was noticeably lower, although still very high.

**Figure 6.**
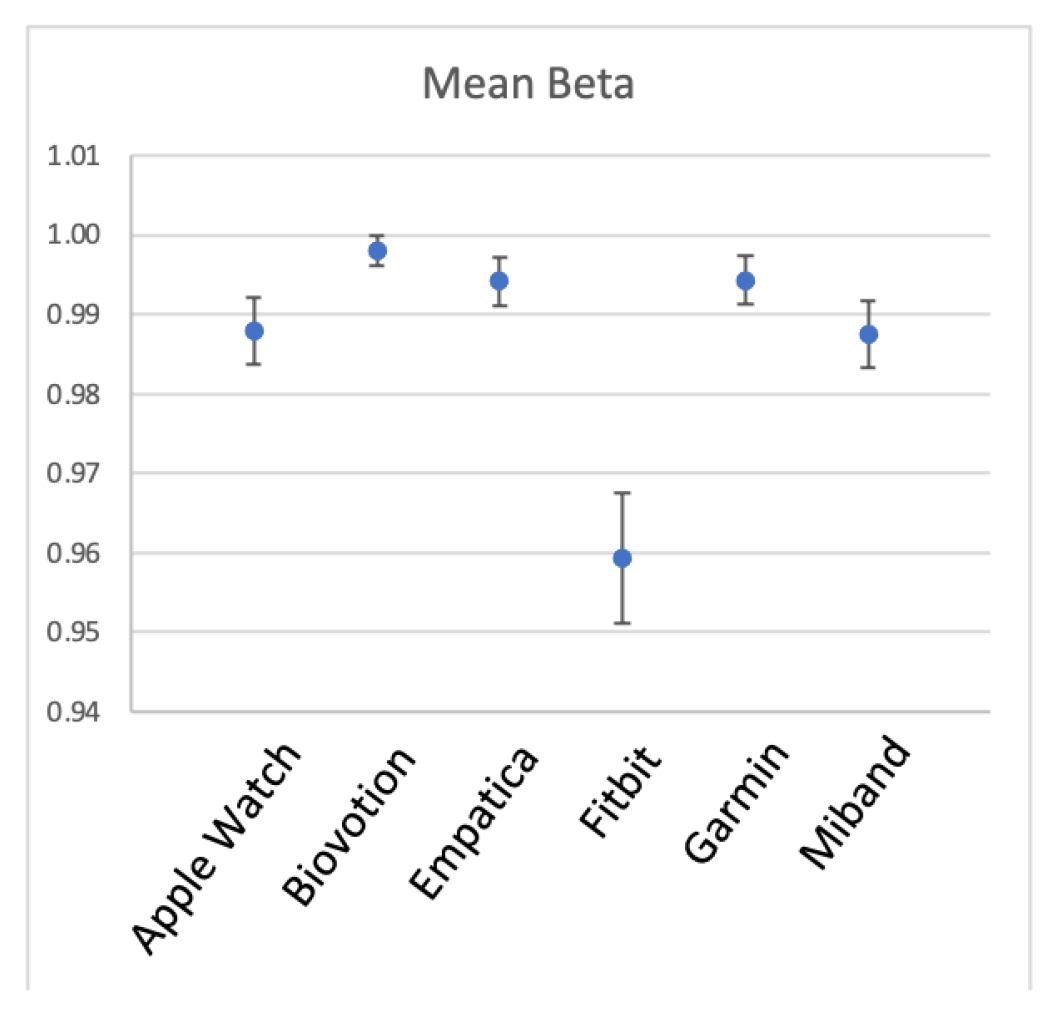
Within-participant reliability of HR for the 6 brands of wearables devices. Error bars represent 1 SD.

**Table 3.** *Within-participant reliability of HR for the 6 wearable sensors, listed in alphabetic order*.

| Device | Mean Beta | SD of Beta across iterations |
| --- | --- | --- |
| Apple Watch | 0.988 | 0.008 |
| Biovotion | 0.998 | 0.004 |
| Empatica | 0.994 | 0.006 |
| Fitbit | 0.959 | 0.016 |
| Garmin | 0.994 | 0.006 |
| Miband | 0.988 | 0.009 |

Figure 7 and Table 4 show within-participant reliability for the 5 activities. It was also very high across the activities, yet transitioning was noticeably less consistent than the other activities.

**Figure 7.**
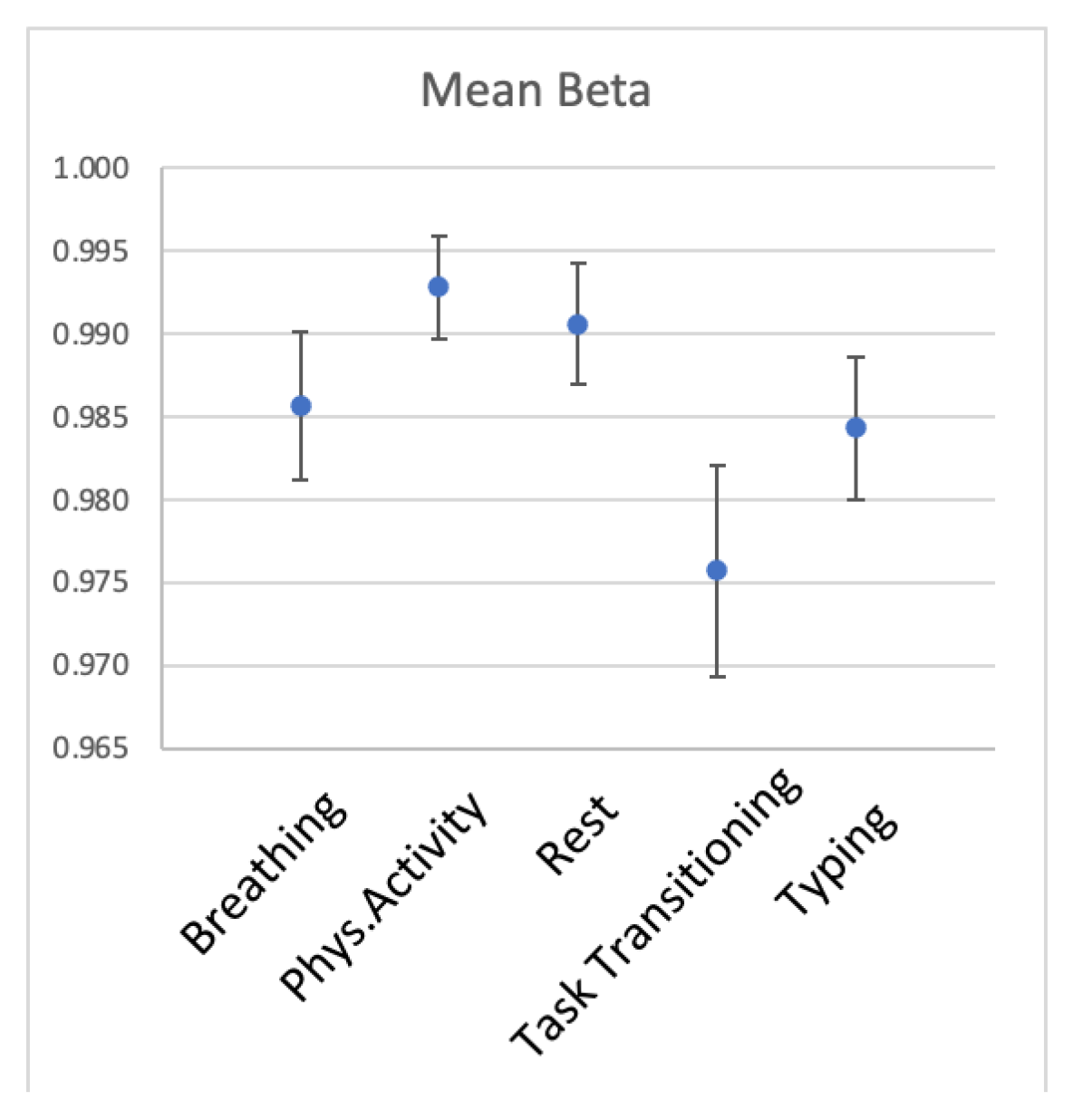
Within-participant reliability of HR for the 5 types of activity. Error bars represent 1 SD.

**Table 4.** *Within-participant reliability of HR for the 5 types of activity*.

| Activity | Mean Beta | SD of Beta across iterations |
| --- | --- | --- |
| Breathing | 0.986 | 0.009 |
| Phys.Activity | 0.993 | 0.006 |
| Rest | 0.991 | 0.007 |
| Transitioning | 0.976 | 0.013 |
| Typing | 0.984 | 0.009 |

### Discussion

We explored the within-and between-participant reliability of heart rate measured with 6 wrist-worn devices during 5 activities. This demonstration showed that measurement fidelity can be estimated without referencing any benchmark device, from the data of a single sensor. We observed noticeable differences in between-participant reliability for the six brands of wearable sensors, and for the different levels of activity participants engaged in. With regard to the two components of measurement fidelity – reliability and validity – the data complied with textbook expectation, showing that high reliability is a necessary, yet not sufficient condition for validity. It showed that validity cannot be inferred from reliability, and that validation of a device is a necessary first step to ensure measurement fidelity under ideal (laboratory) conditions. Yet, as the analysis of different activity levels showed, even once an acceptable level of validity is established under resting conditions, a wearable device can produce measurement of suboptimal reliability under more active everyday conditions.

Within-participant reliability was very high across devices and activity levels. Heart rate is a great example of a measurement that is highly consistent within-participant (high within-participant reliability), but not always acceptable for distinguishing between participants (moderate between-participant reliability). This most probably reflects the nature of heart rate, which has stable and quite narrow limits for a given person, especially during wakeful time. In our next example (Study 2) we examined a less constrained measure — heart rate variability — in order to see how within-and between-person reliability is manifested in this measure.

## Study 2: Reliability of Biostrap during sleep and during wakeful time

In this study we tested a commercially available Biostrap wristband sensor for both between-person and within-person reliability of HR and HRV. In our treatment of within-person reliability, we focused on comparing two diurnal states of the user: wakefulness and sleep. Sleep corresponds to time passing with little to no change in the external environment and fewer physiological changes than during wakeful periods (e.g., relatively little physical effort, no eating or talking, relatively little stress and mental effort). We hypothesized that sleep periods would produce less variable and therefore more reliable biometric recordings.

Ten participants wore a Biostrap device continuously for one week. They were instructed to wear the device on their wrist at all times, except when charging the device (about 1 hour daily) and when taking a bath or shower.

### Method

#### Participants

Ten participants (1 male) were recruited through Reservax (https://www.reservax.com), an online recruitment platform for behavioral studies. The inclusion criteria were: Participants at least 18 years of age, without known heart problems or disease, in generally good health, and fluent in written and spoken English. All participants provided informed consent prior to participation. Participants were paid a maximum of $100 CAD for participation, based on their compliance with study procedures. All 10 participants received the full payment.

#### Apparatus

The Biostrap wristband is a commercially available PPG sensor of heart rate (https://biostrap.com). Biostrap (formerly Wavelet) has been validated against clinical-grade wearable devices and ECG (Dur et al., 2018; Jarchi et al., 2018; Steinberg, Yuceege, Mutlu, Korkmaz, van Mourik, et al., 2017). The device uses long wavelength light (red) to detect pulse. Automatic sampling is performed once in every 5 minutes (in enhanced mode), each recording lasting for 45 seconds at 43 Hz frequency. The raw data are stored on the sensor’s internal memory, then transmitted to a smartphone app via Bluetooth connection, and then to the Biostrap server where the data is processed.

The output provided by Biostrap includes: heart rate in beats per minute (BPM), heart rate variability (HRV) indexed as the root mean square difference between successive heartbeats (rMSSD), oxygen saturation, and respiration rate. This information is provided for each sampled measurement, which can be as frequent as once in every 5 minutes. The sensor also includes an accelerometer, which provides information on the number of steps completed by the wearer. Based on a combination of these metrics, sleep onset and offset are detected.

The commercial Biostrap smartphone app ordinarily shows the user their heart rate and heart rate variability, number of steps, and a sleep score on the app’s home screen (these metrics are shown by default). It also indicates the battery status and the last time the data were synchronized with the app. In this study the app was blinded to participants, so that it was unable to display any biometrics; only the battery status was visible to them. Ecological momentary assessments were delivered using the Ipromptu smartphone app (http://www.ipromptu.net)

#### Procedure

Invited participants arrived at the lab in the Department of Psychology at UBC for an introductory session, where they were introduced to the Biostrap device, provided personal demographic information, and completed questionnaires on emotional, self-control, and personality traits (which are not reported here).

Each participant received a fully-charged Biostrap wristband to wear for the duration of the study along with a charging plate. The Biostrap app was installed on participants’ smartphones and they were instructed on the use the sensor and how to ensure the data were synchronized regularly. Instructions to participants emphasized that they were to wear the device at all times, including times of exercise and sleep, except for when charging the device or taking a shower or bath. Participants wore the Biostrap continuously for 8-11 days

In addition, participants were asked to track their emotional state using an ecological momentary assessment (EMA) approach. Ipromptu app (http://www.ipromptu.net) was used to deliver short surveys 6 times a day, at random times between 8 am and 8 pm. If not responded, a prompt repeated twice, with 15-minute intervals, and was available for response for several hours. Participants were instructed to respond to at least 1 and as many prompts as they could. On each prompt, a 8-question survey asked participants to rate, on a scale from 1 to 10, how happy / energetic / nervous / afraid / irritable / angry they are and how much pain and discomfort they were feeling, in random order.

Participants returned to the lab at least 8 days after their introductory meeting to conclude the study. They returned the Biostrap devices, were debriefed about the purpose of the study, and paid for their participation.

The study procedures were approved by the institutional Research Ethics Board (approval number H19-01197). Data, materials, and analysis code for this study are available at https://zenodo.org/badge/latestdoi/520639317.

### Results

#### Data processing

The heart-rate measurements consisted of raw PPG waveforms, which were processed by Biostrap’s algorithms in their servers (Dur et al., 2018). The data presented here were based on the aggregated metrics provided by the Biostrap for each successful sample, which included beats per minute (BPM) and heart rate variability (HRV), calculated as the root mean square difference between successive heartbeats (rMSSD). Hereafter we will refer to HRV for simplicity, instead of rMSSD. These heart-rate measures were further screened for artifacts and anomalies in two steps. First, all 0 values were removed (affecting 0% of BPM and an average of 19.26% of HRV samples across all participants). Second, values exceeding each participant’s mean by more than 2 standard deviations over the whole observation period were removed (affecting 2.84% of BPM, and 2.19% of HRV across all participants).

Participants’ state (asleep vs. awake) was established using the heart-rate indices in the following way. We found periods of at least 2 hours in duration when BPM samples were successfully recorded at least every 15 minutes. We then chose the longest such period on each day and assumed that it corresponded to sleep. Although time of day was not a criterion for determining sleep periods, all the sleep periods established in this way happened to occur between 9 pm and 11 am. These criteria allowed us to detect at least 5 periods of sleep for 8 of 10 participants. We recognize that these criteria do not guarantee that participants were awake at all other times, and as such, that this potentially biases awake observations to be appear to be more similar to sleep periods. But to anticipate the results, the density and reliability of HR and HRV assessment during sleep periods defined in this way were greater by orders of magnitude than they were during the defined wakeful times.

Heart-rate data was successfully recorded for only 2 sleep periods for one participant and only 1 sleep period for another, and so their data were not included in the analyses. Days with only one HRV sample during wakeful times (5 periods across participants) were also excluded from the analyses. All participants had more than one HRV sample during sleep. These exclusions left us with 8 participants tracked continuously for 5 to 11 days, and a total of 6840 samples for BPM and 5530 samples for HRV.

#### Descriptive statistics

Figure 8 shows the frequency of successful heart-rate samples for BPM and HRV made during wakeful and sleeping periods. The pattern of these two variables was generally consistent across participants. The mean number of BPM samples acquired for waking periods was 19.42 (SD = 13.52), and the mean number of sleep samples was 74.6 (SD = 24.43). The mean number of HRV wakeful samples was 13.65 (SD = 9.63) and the mean of sleep samples was 69.70 (SD = 22.63).

**Figure 8.**
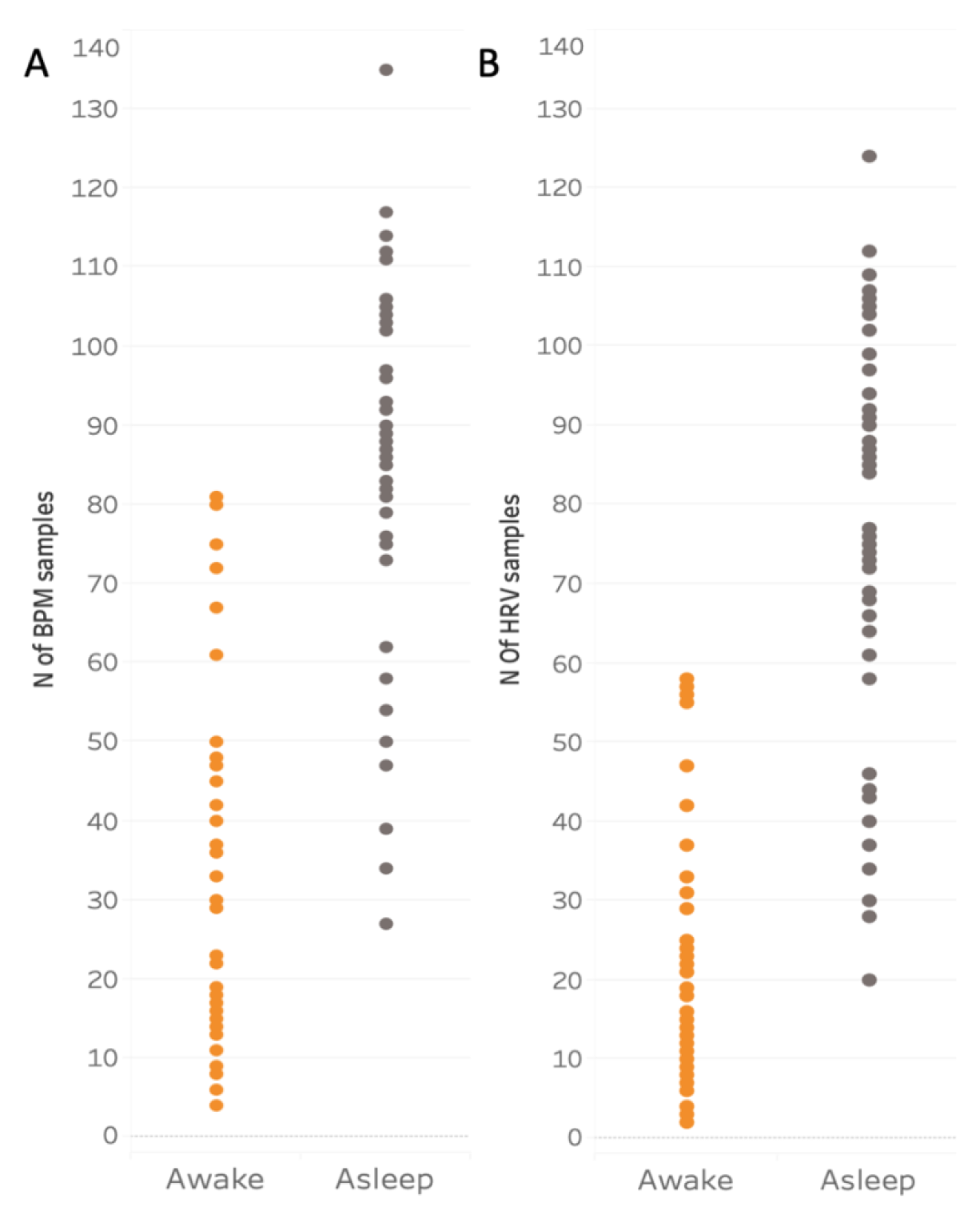
Average number of successful measurements of BPM and HRV per participant per day during wakefulness and sleep. **Panel A** shows average number of successful measurements of BPM, **panel B** shows the same data for HRV per day and per participant during wakefulness (orange) and sleep (grey).

Figure 9 shows the mean BPM and HRV for each participant, separately for wakefulness and sleep. The figure shows that there are pronounced individual differences in both biometrics, with some participants having consistently higher HRV or BPM than others. The variability of the wakeful measurements is also visibly larger than the variability of sleep measurements. These observations were confirmed by the following analyses.

**Figure 9.**
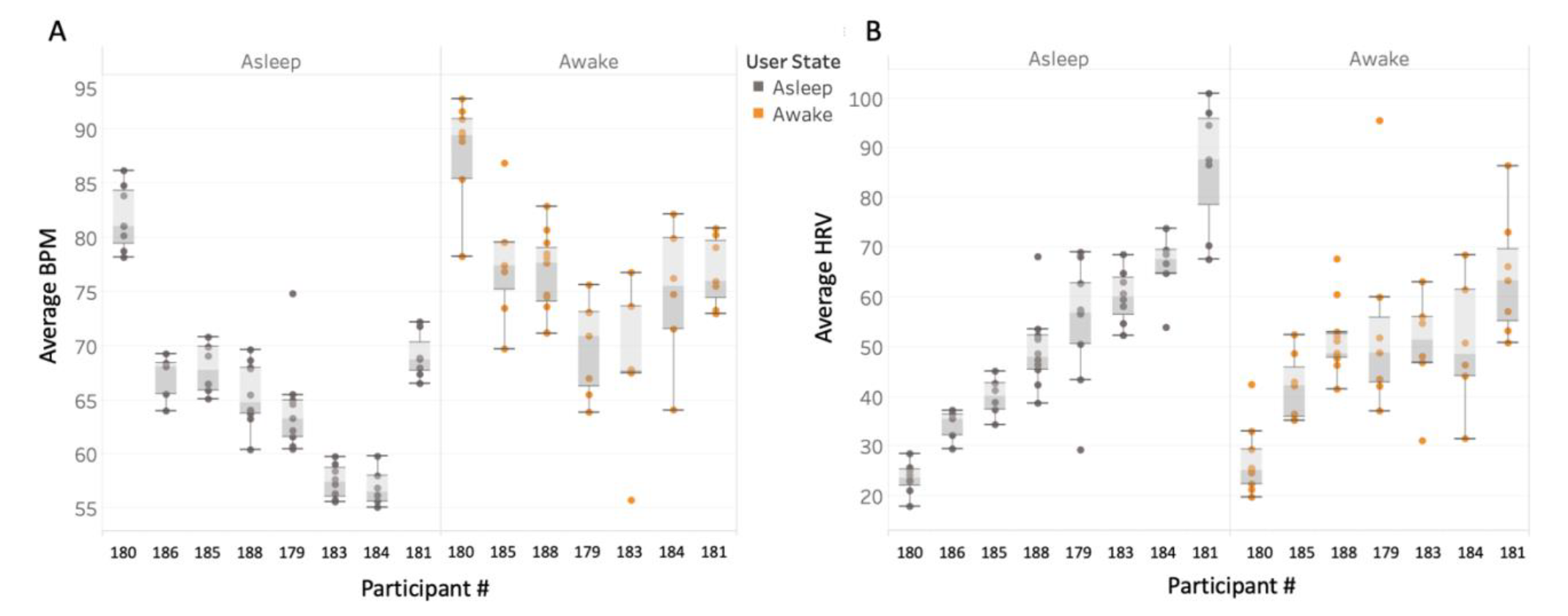
Average HR and HRV per participant during sleep and wakefulness. **Panel A**: The mean BPM for each participant, separated for sleep and wakefulness. **Panel B**: The mean HRV for each participant, separated for sleep and wakefulness. Participants are rank ordered in each panel based on their HRV during sleep. Participant 186 had no biometric recordings for wakeful time.

#### Between participant reliability

A one-way random single-measure ICC(1,1) was computed for the 8 participants’ mean BPM and HRV, separately for wakeful and sleep period samples. ICC values were consistently higher for sleep periods than for wakeful periods. This was true for both BPM values (sleep: *ICC(1,1)* = .89.6, 95%CI [.79 .97], *p* < .001; wakefulness: *ICC(1,1)* = .55, 95%CI [.34 .83], *p* < .001) and for HRV values (sleep: *ICC(1,1)* = .84, 95%CI [.70 .95], *p* < .001; wakeful: *ICC(1,1)* = .39, 95%CI [.19 .73], *p* < .001). These high ICC values for sleep, along with only moderate ICC values for wakefulness, imply that individual differences in heart rate and heart rate variability can be measured more reliably with a commercial PPG sensor during sleep than wakefulness.

These data suggest that BPM and HRV measured through a commercial wearable device are relatively stable between people, meaning that a person whose BPM or HRV is higher than other people’s on one day/night is likely to have BPM or HRV higher than other people on any other day/night.

#### Within-participant reliability

Within-participant reliability was assessed using a split-half approach and mixed model regression. We compare two methods of splitting the data: time-sensitive (split into odd and even samples, by order of measurement) and random (dividing the datapoints into two samples randomly, so that a sample from early in the day is equally likely to be paired with a sample from later or earlier in the day). For both methods, a mixed model regression is then computed predicting one estimate of the biometric (e.g., average of the odd datapoints) from the other estimate (e.g., average of the even datapoints), with participant as the random factor (see formula 3), and Satterthwaite’s correction for the degrees of freedom.

The time-sensitive method resulted in estimates of the within-participant reliability of BPM and HRV illustrated in Figure 10. Panel A shows that reliability of BPM was very high for the sleep and wakeful periods alike. The effect of predictor BPM was highly significant in both models, *beta* = 0.99, *t*(9.17) = 91.8, *p* < .001 and *beta* = 0.82, *t*(18.8) = 7.91, *p* < .001, respectively. Panel B shows that the fit between predictor and criterion for HRV was also generally high during sleep, *beta* = 0.96, *t*(57) = 51.18, *p* < .001, but not during wakefulness, *beta* = .097, *t*(47.26) = 0.92, *p* = .36.

**Figure 10.**
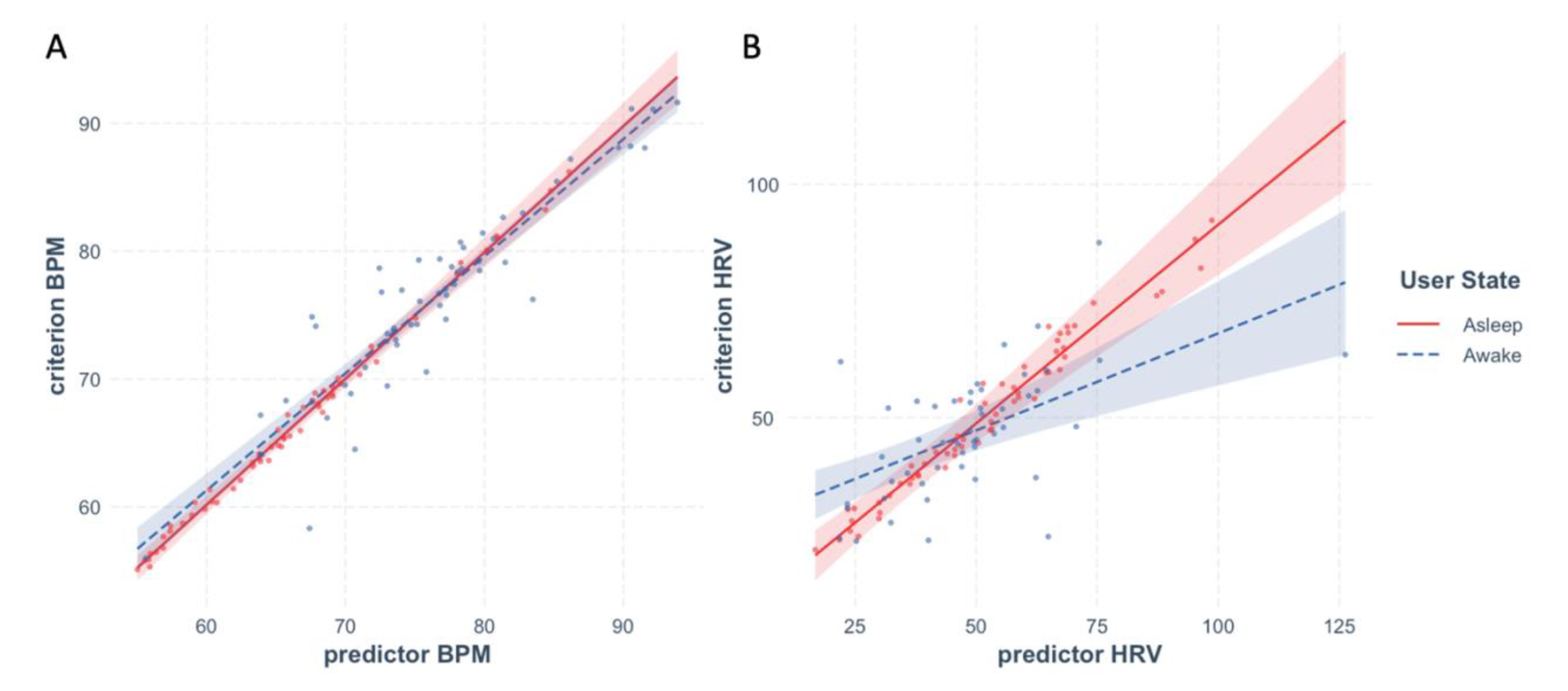
Within-participant reliability of HR and HRV during sleep and wakefulness estimated with the time-sensitive method. **Panel A** shows within-participant reliability of BPM, **panel B** of HRV. Data recorded during sleep is shown in red, during wakefulness in blue. Shaded area represents 95% CI, dots represent partial residuals.

Random splitting into the subsamples, as mentioned above, can result in extraordinarily low or high estimate of reliability. Therefore, we performed the split 1000 times, computing mixed-model regression each time. Figure 11 shows distributions of the resulting *beta* values for HR and HRV during sleep and wakefulness. We estimated reliability as the average beta across the 1000 iterations.

**Figure 11.**
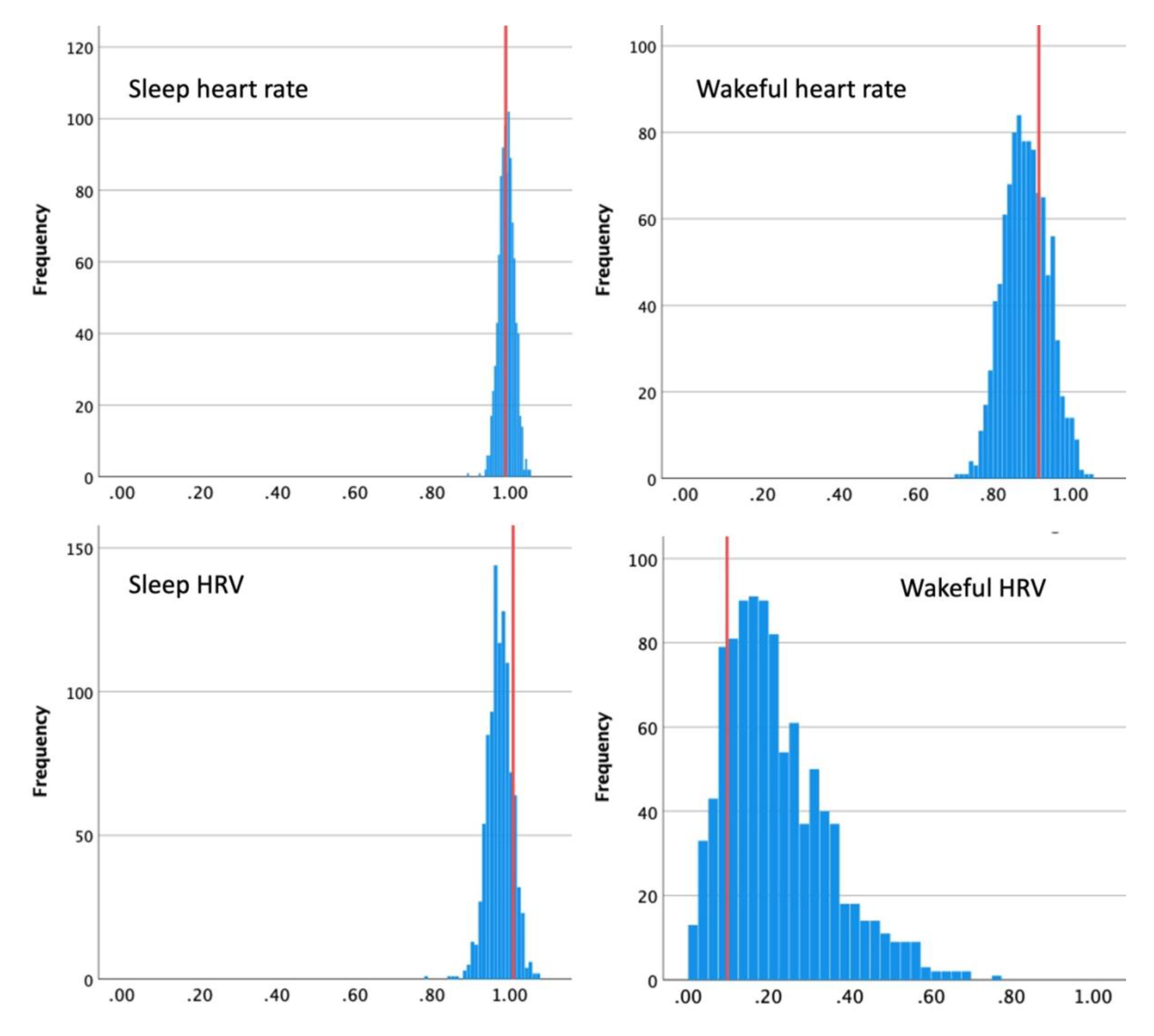
Within-participant reliability of HR and HRV during sleep and wakefulness estimated with the random approach. Each graph shows a distribution of betas for respective cardiac biometric, with red lines showing beta estimated with the time-sensitive method.

For BPM, random method produced reliability estimates that were very close to those resulting from the time-sensitive approach, if slightly lower during wakeful time, *M*_beta_sleep_ = .99, *SD* = .02, *M*_beta_wakeful_ = .89, *SD* = .058. For HRV during sleep, reliability from the random method was slightly lower than from the time-sensitive approach, *M*_beta_sleep_ = .98, *SD* = .03, supporting our assumptions. However, for wakeful HRV random approach resulted in somewhat higher estimated reliability than that produced by time-sensitive method, *M*_beta_wakeful_ = .22, *SD* = .13.

To summarize, the two methods of estimating within-participant reliability revealed that both BPM and HRV were highly reliable during sleep, BPM was also very reliable during wakeful time, yet reliability of HRV during wakeful time was drastically lower. The two methods diverged in assessment of the latter, and not in the predicted direction, with the time-sensitive method yielding much lower estimate of reliability. Notice, however, that the number of datapoints obtained for HRV during wakeful hours was much lower than for sleep-time HRV or for BPM during wakefulness (see descriptive statistics above). Therefore, the amount of time separating successive datapoints must have been particularly long for wakeful HRV, likely exceeding the period during which we would expect such a volatile measure as is HRV to be stable. The fact that the range of reliability estimates obtained with the random method was extremely wide (0.2 – 1) supports this reasoning.

### Interim discussion

We have demonstrated how between-person and within-person reliability can be estimated in data readily available from a commercial wearable sensor of cardiac biometrics. For the particular device tested here, between-person reliability as assessed with ICC was excellent for sleep-time HR and HRV, but only moderate for wakeful biometrics. Within-person reliability, assessed using split-half and mixed model regression approach, was near-perfect for HR during sleep as well as wakeful time, but HRV was only reliable within-person during sleep, not during wakefulness.

It is worth noting that periods of lower reliability in this study coincided with periods in which fewer datapoints were obtained (lower measurement density). This could be taken to suggest that increased measurement density contributes to greater reliability. Yet this coincidence in the present study should not be interpreted too strongly, since heart rate during wakeful times had a relatively high within-participant reliability even in the face of relatively fewer data points (lower measurement density). Further studies should investigate whether and how much measurement density contributes to higher reliability.

One immediate consequence of the compromised reliability of a measure is the reduced ability to detect its relationships with other variables (Spearman, 1904). We demonstrate this in what follows by testing whether BPM and HRV can be predicted from subjectively reported emotional states of the participants. Multiple laboratory studies showed that stress and cardiac biomarkers are strongly associated, and this relationship was recently replicated with wearable sensors (Coutts et al., 2020; Hovsepian et al., 2015). We had no prior hypothesis as to which of the biomarkers (BPM, HRV) would produce stronger association if they were measured with equal fidelity.

### Correlations between biomarkers and subjective emotion

To analyze subjective emotions captured with the EMA, we averaged the 4 negative emotions on each prompt (irritable, afraid, nervous, angry), and the 2 positive emotions (happy, energetic). We then averaged responses to all the prompts within one day, which resulted in two scores per day: one for positive and one for negative emotions.

We then used these two scores (negative and positive emotions) as predictors in mixed model regressions with participants as random factor. We first tested wakeful BPM (and wakeful HRV in separate analyses) on the concurrent day as the dependent variables. Then we tested the same models on sleep BPM (and sleep HRV in separate analyses), either on the preceding night (two models) or following night (two more models). This meant that six models were tested in all, prompting us to use Bonferroni corrections to test for significance. The strongest relationship in all of these involved a negative one between sleep-BPM and lower negative mood reports on the subsequent reporting day, *beta* = −2.11, *t*(40.25) = −2.745, *p* = .054 (Bonferroni corrected). This meant that when a participant experienced higher sleep-BPM they were less likely to report negative emotions on the following day; when they experienced lower sleep-BMP they were more likely to report negative emotions the next day. Wakeful-BPM was not reliably associated with either positive or negative emotions, *ps* > .5 (uncorrected).

For HRV, the strongest effect was also one where relatively higher sleep-HRV on a given night predicted greater negative emotions on the subsequent day, *beta* = 4.45, *t*(40.5) = 1.79, *p* = .081 (uncorrected). Wakeful-HRV was not associated with either positive or negative emotions, *ps* > .2 (uncorrected).

**Figure 12.**
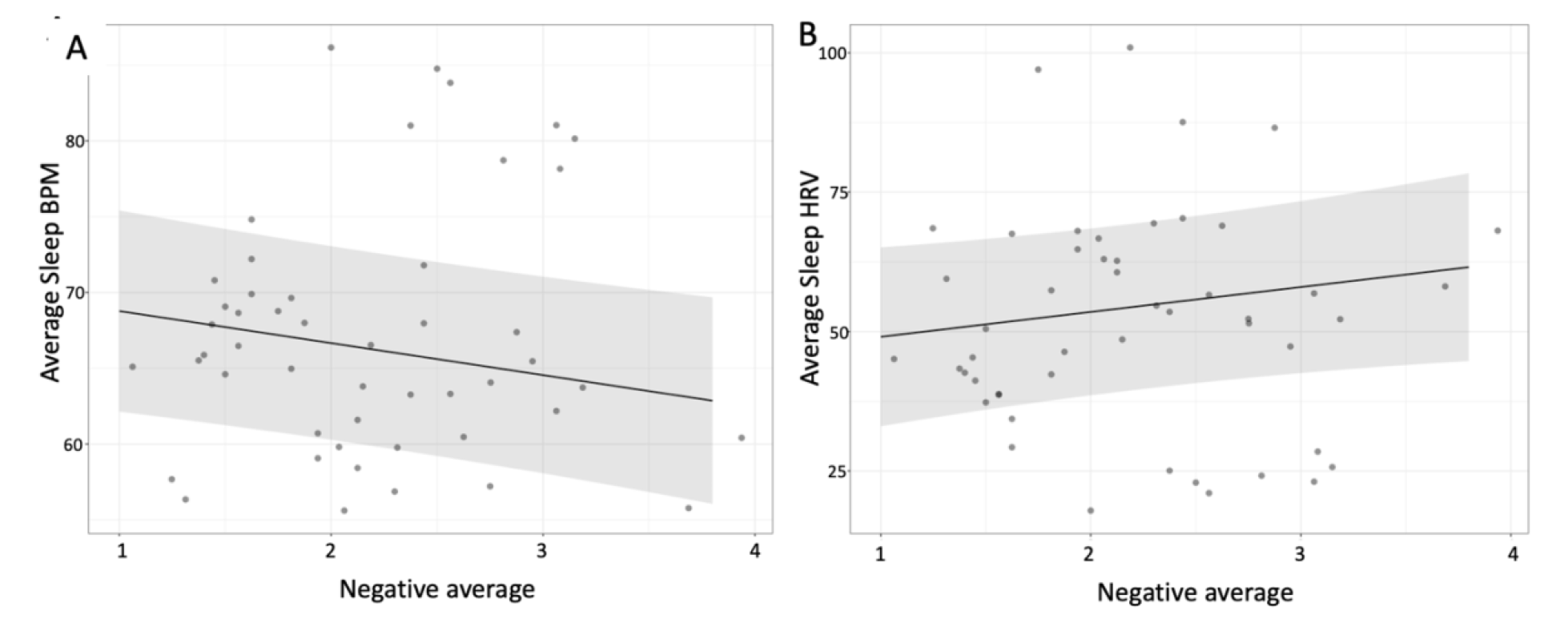
Association between negative emotions and sleep HR(V). Predicting sleep-BPM (left panel) and sleep-HRV (right panel) from negative emotions on a subsequent day. Grey area represents 95% CI.

### Interim discussion

Our attempt to test the association between mood during a day with cardiac biometrics concurrently (wakeful), on preceding or following night, revealed a predictive relationship between night-time BPM and mood on subsequent day. We cannot tell from this study whether the difference between the results for BPM and HRV stems from difference in reliability only or for other reasons unrelated to measurement fidelity. However, the difference between wakeful and sleep BPM in predicting daytime mood is surprising, given that both mood and cardiac biometrics change rapidly, so the closer in time they are measured the higher we would assume the association to be. It is therefore plausible that the association between daytime BPM and mood was compromised by the lower reliability of daytime measurement.

## General Discussion

The validity of data from wearable sensors is now thought to be quite good (Barrios et al., 2019; Dur et al., 2018; Hernando et al., 2018; Kinnunen et al., 2020; Menghini et al., 2019; Steinberg, Yuceege, Mutlu, Korkmaz, van Mourik, et al., 2017). However, the reliability of the biometrics captured by these devices in daily life has so far been assumed to be high, but it has rarely been tested systematically. Interestingly, recent reports of using wearable sensors of HR & HRV outside of laboratory or clinical settings have revealed that validity of data from wearable sensors (i.e., correlations with a criterion, usually a medical-grade wearable device) in the conditions of everyday life is lower than expected from laboratory studies where participants are typically at quiet rest (Galarnyk et al., 2019; Sneddon & Carlin, 2019). The finding by Van Voorhees et al. (2022) that the reliability of a wearable (Empatica E4) HRV measurement across 24 hours was unacceptably low offers an account of low validity of wearable sensors outside of a lab. Our examination of wearable data reliability in the present study also suggests that day-time measurements of HR and HRV are highly unreliable. Yet rather than be discouraged by these data, we suggest focusing on *how* wearable sensors can be used to deliver data that is reliable. This can be done by assessing the sensor’s reliability in different situations, such as we did here for periods of sleep and wakefulness. This approach has been taken previously in the fields of movement science and athletics, where there was a growing awareness of the importance of testing the reliability of many forms of sensing technology in varying environmental and situational contexts (Evenson & Spade, 2020; Kobsar et al., 2020; Kooiman et al., 2015; Straiton et al., 2018).

Here we applied the theory of reliability developed for psychological questionnaires to physiological measurements obtained with a wearable device. In doing so, it is important to keep in mind the research goals and questions. If measurements are performed for comparisons between persons, between-participant reliability should be assessed, e.g., using ICC. If, however, the aim of measurement is to detect different states within the same person, within-person reliability should be estimated, e.g., using a combination of split-half and mixed-model approach. While estimating between-participant reliability is straightforward, estimating within-person reliability, without additional devices and measurements, must take into account the time-sensitive nature of the measurements being made. In the case of HR and HRV, the passage of time is a critical variable. Other physiological measurements will likely have similar considerations that are specific to the type of measurement being made.

We applied this approach to commercially available PPG sensors of cardiac biometrics, showing how between-and within-person reliability could be estimated from open-source data. The results showed that both between-and within-participant reliability of heart rate measurement varies for the different brands of wearables. Across different brands, it also varies for the different levels of user’s activity. This suggests that even the most reliable of the sensors tested (Apple Watch, Biovotion) may produce more or less reliable measurement in different circumstances.

Focusing on the Biostrap wearable sensor with or own data, we found that the between participant reliability of HR and HRV was excellent during sleep (ICC > .75), but only fair during wakefulness (ICC [.4 .6]). Within-participant reliability of HRV was also found to be higher during sleep than during wakefulness. Finally, we found that correlations of HR and HRV with a second variable – in our case, subjectively reported mood – were stronger for the most reliable metric, sleep-time BPM. Taken as a whole, the present data suggests that the wearable sensor we used (Biostrap) provides data that is highly reliable during sleep, and less so during wakefulness.

The most popular testing of wearable devices has focused so far on measuring their validity during different levels of physical activity (Barrios et al., 2019; Thomson et al., 2019)). This is not surprising given that the early vision for the application of wearable devices was to regulate exercise load for performance optimisation. This is still the primary use of many wearable devices today (Almeida et al., 2019; Thomson et al., 2019). These studies have shown that that some measurements cannot be taken reliably during physical activity (Almeida et al., 2019). At the same time, other studies have shown that the application of wearables is not limited to detecting acute events (such as exercise load or acute stress), but that they can be useful in indexing slower fluctuations in the user’s state, such as overall physical shape or allostatic stress, which affect one’s long-term health and wellbeing (Chuang et al., 2015; Koskimäki et al., 2019). This development opens up a unique research opportunity to measure psychophysiology longitudinally, across a variety of real-world contexts and extensive time periods (Kleckner et al., 2021). And this purpose might be best achieved with measurements taken during acute events or during recovery after those events. Indeed, a large body of studies have begun to investigate stress by focusing on recovery after stress, rather than on what is going on at the moment of acute stress (Allen et al., 2014). The hope is that by systematically quantifying measurement reliability in different circumstances, researchers will eventually be able to make informed choices about specific wearable devices and measurement procedures that meet their research goals.

## Footnotes

1 In practice values close to −1 are not usually observed and would suggest existence of a strong source of systematic variance in the measurement. Negative numbers near 0 suggest low reliability.

